# Microdomes: Micro-engineered Microdome arrays enable standardised and shear-free 3D biology with full-spectrum optical imaging compatibility

**DOI:** 10.64898/2026.09.10.750786

**Authors:** Mei Shan Cheam, Saburnisha Binte Mohamad Raffi, Hui Ting Ong, Sandra E. Ghayad, Marion Le Grand, Eddy Pasquier, Phoebe Dunbabin, Kate Poole, Jasmine Chin Fei Li, Liu Jun, Esra Karatas, Natalie Lim Sheng Jie, Joanne Lim Tze Chin, Modhura Ganguly, Veronique Angeli, Yusuke Toyama, Gianluca Grenci, Anne Beghin

## Abstract

Three-dimensional cellular models such as organoids and spheroids hold major promise for developmental biology, disease modelling and precision medicine, yet their large-scale production and analysis remain constrained by handling-induced shear stress, sample fragility, positional instability and limited compatibility with advanced imaging workflows. Here, we introduce Microdomes, dense arrays of open-top, dome-shaped microcavities. Each cavity holds a single specimen in its own miniature aquarium, accessed through a narrow apical opening that admits cells, medium, matrix and staining reagents while shielding it from the shear generated during routine pipetting. Each Microdomes chip accommodates more than 100 spheroids or organoids, supporting diverse cell types, co-culture formats and both matrix-free and matrix-embedded culture. All specimens are retained through prolonged culture, repeated medium exchange, fixation and immunostaining. Because every specimen occupies a fixed, addressable position against a thin transparent film, the same spheroid can be relocated and re-imaged over weeks of culture and across microscopes, from array-wide overviews to subcellular details, without transfer, embedding or other perturbation. Microdomes also support AI-based automated segmentation, from whole-spheroid outlines to individual nuclei in 3D, using common image analysis software. Microdomes thereby turn each array into a self-contained quality control unit, providing specimen-level traceability that organoid production pipelines currently lack.

## Introduction

Three-dimensional (3D) biological systems, including organoids, spheroids and engineered microtissues, have transformed our ability to model development, physiology, disease and therapeutic responses. Their broader adoption, however, remains constrained by limitations in culture standardisation, sample handling and optical interrogation. Many existing formats provide limited control over specimen size and position, complicate high-throughput workflows, and frequently restrict optical access, limiting the use of advanced imaging modalities. Repeated medium exchange and post-culture processing can also displace, deform or result in the loss of delicate structures, causing experimental attrition and potentially biasing analyses toward specimens that are more readily retained. A practical platform for 3D biology must therefore combine reproducible specimen organisation with minimal handling-induced perturbation, reliable sample retention and broad optical accessibility.

Embedding specimens within domes or layers of basement membrane extract, most commonly Matrigel, remains one of the simplest and widely used approaches to 3D culture, requiring no specialised device. This format underpinned seminal organoid culture protocols and remains in routine use today^1,2^. However, specimens are distributed unpredictably within bulk matrices, resulting in variable imaging depths, complicating longitudinal tracking and scalable imaging. Micro-engineered culture systems address some of these limitations by imposing defined geometries and organising specimens at predetermined locations. Commercial microwell platforms, including InSphero® Gri3D® Hydrogel Microcavity Plates^3^, AggreWell™ Microwell Plates^4^ and Corning® Elplasia® Plates, support the parallel formation of relatively uniform 3D tissues within ordered arrays, facilitating high-throughput culture, imaging and screening. Some platforms incorporate features intended to minimise specimen disturbance during liquid handling, such as the segregated pipetting port of the Gri3D Hydrogel Microcavity Plate. Nevertheless, their geometries can constrain high-resolution in-plate imaging, while the effects of handling-induced fluid shear during routine liquid exchange remain poorly defined^3,4^. Recent platforms have extended these capabilities through permeable nanofibrous microwells, as exemplified by the Uniform and Mature organoid culture platform (UniMat)^5^, and through image-addressable microwells that enable longitudinal phenotyping and selective organoid retrieval^6^. Droplet-based microfluidic systems similarly offer highly parallelised and compartmentalised 3D culture^7–9^. Nevertheless, these platforms generally address specific aspects of 3D culture, such as uniform formation, high throughput or selective retrieval, rather than integrating these capabilities with direct high-resolution multimodal imaging and protection from handling-induced mechanical perturbation.

To overcome these longstanding bottlenecks, we developed Microdomes, a new generation of micro-engineered array designed to integrate standardised 3D culture with optical accessibility and shear-free handling. Each microdome functions as a miniature aquarium that holds a single 3D specimen at a predefined and permanently addressable position, shields it from handling-induced flow, and permits routine medium exchange and downstream processing through its open top. The thin, transparent construction of the device further enables specimens to be interrogated using a broad range of imaging modalities, including brightfield and phase-contrast microscopy, laser-scanning and spinning-disk confocal microscopy, and two-photon microscopy. The platform accommodates extracellular matrix (ECM)-embedded, ECM-free and hybrid culture formats and can be incorporated into diverse experimental configurations, including multi-well plates, for scalable experimentation.

The Microdomes are inspired by the JeWells we previously developed for single-objective SPIM (soSPIM)^10^, but have been re-engineered to ensure full compatibility with conventional 3D imaging modalities, larger biological specimens, and routine long-term 3D culture. An isotropic hemisphere replaces the pyramidal cavity, avoiding sharp edges that may bias cell behaviour and introduce optical aberrations, and the enlarged dimensions accommodate a wide range of seeding densities, keep specimens within the working distance of most objectives, and allow efficient cell loading while preventing specimen escape.

Here, we describe the design, fabrication and validation of Microdomes as a versatile platform for 3D culture, imaging and analysis. Using diverse cell types, we demonstrate compatibility with multiple culture configurations. We quantify the negligible shear stress experienced by specimens during routine liquid handling and demonstrate complete sample retention during long-term culture and throughout fixation and staining. Finally, we show that individual specimens can be longitudinally tracked and interrogated across multiple microscopy platforms, enabling correlative imaging from array-scale observations to high-resolution 3D analysis. Together, these capabilities position Microdomes as an accessible bridge between scalable 3D culture and advanced imaging workflows.

## Results

### Geometric design and fabrication process of the Microdomes

The Microdomes used here are produced by a soft-lithography process previously described^10–12^. Each Microdome is a hemispherical cavity of 600 µm diameter with a circular top opening of 100 µm diameter, extended into a cylindrical "chimney"-like entry port of 20 µm height (**Fig. 1a**, **right)**; the cavities are arranged as a square array at 800 µm pitch within a 10.5 × 10.5 mm^2^ footprint, giving 144 cavities per chip (**Fig. 1a**, **left)**.

**Fig. 1.**
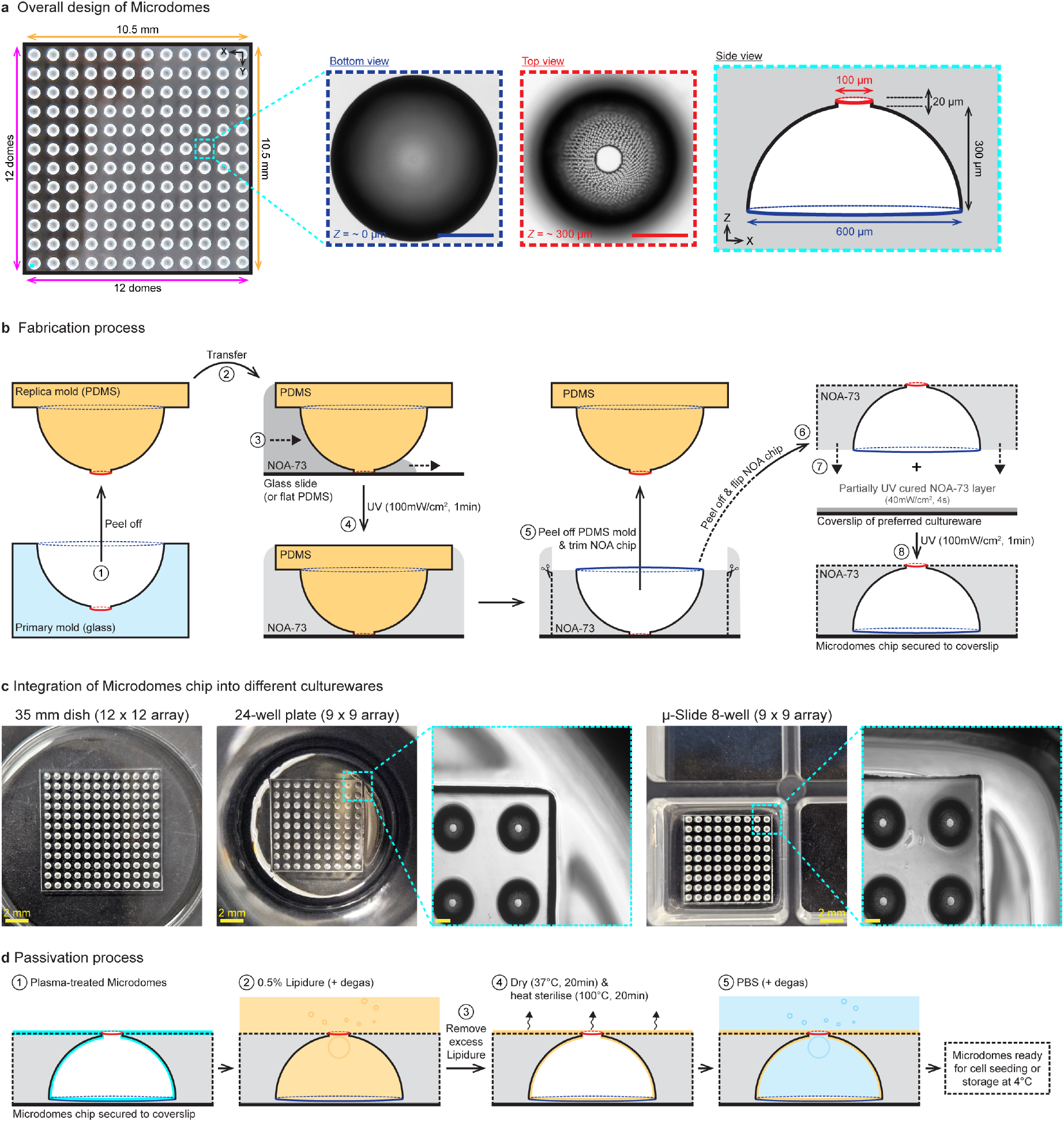
Design and fabrication of the Microdomes. **a.** Geometric design of the Microdomes. Left: top view of the Microdomes chip, a 12 × 12 array of hemispherical cavities at 800 µm pitch within a 10.5 × 10.5 mm footprint. Middle: brightfield images of a single dome acquired at its base (z ≈ 0 µm) and top (z ≈ 300 µm). Right: cross-sectional schematic of a dome, 600 µm in diameter and 300 µm in height, with a cylindrical opening of 100 µm in diameter and 20 µm in height. **b.** Conceptual scheme of the Microdomes chip fabrication process. (1) A PDMS mold is cast as a replica of a glass primary mold. (2) The PDMS mold is placed over a glass slide or flat cut of PDMS, and (3) the intervening space is filled by capillarity with the UV-curable resin NOA-73 (dotted arrows indicate the direction of liquid progression). (4) UV exposure cures the resin into a solid film, which is (5, 6) trimmed and flipped, then (7) placed onto a coverslip pre-coated with a thin, partially cured NOA-73 layer. (8) A final UV exposure secures the Microdomes chip to the coverslip. **c.** Integration of Microdomes chip into different commercial culturewares, including cell culture microscopy dish, well-plate and chamber slide. **d.** Surface passivation process of integrated Microdomes chip. (1) Surfaces are plasma-treated before (2) addition of 0.5 % Lipidure solution, with degassing to ensure complete surface passivation. (3) Excess Lipidure is removed and (4) the device is fully dried before heat sterilisation. (5) PBS is then added and degassed, leaving the Microdomes ready for cell seeding or storage at 4 °C. Scale bars, 200 µm (a) and 20 µm (c), unless otherwise stated.

The Microdomes chip is produced as a thin film of NOA-73, a biocompatible, UV-curable resin that is optically clear with a refractive index of 1.56 when cured. The film can be bonded to a glass substrate either permanently or reversibly, and accommodates substrates of essentially any shape or size, giving considerable flexibility in the choice of cultureware. Detailed fabrication procedures are provided in our previous publications^10,11^; here we give a simplified but complete description (**Fig. 1b**). A primary glass mold bearing the array of cavities was replicated in PDMS by standard soft lithography. The PDMS replica is cut to its final size (∼10.5 × 10.5 mm^2^) and placed onto a glass slide or a flat PDMS substrate; the cavity formed between the two pieces is filled with the UV-curable resin by capillary action. The resin is cured by UV exposure, and the PDMS mold is then peeled away. The resulting textured film is lifted from this flat substrate, trimmed and inverted onto a coverslip pre-coated with a thin, partially cured NOA-73 layer that acts as a glue; a final UV exposure secures the chip to the coverslip. Manual application of the glue followed by removal of the excess – requiring no spin coater or other specialised coating equipment – gives a bonding layer of approximately 36.7 ± 5.8 µm (*n* = 5 domes sampled at corner and central positions across one chip) **(Supplementary Fig. 1a)** that holds the chip securely while preserving the optical access exploited throughout the results below. Because the chip is a thin solid film, it can be cut to size at the bench and mounted in most common cultureware compatible with high-resolution imaging — microscopy dishes, multi-well plates and chamber slides. A standard chip carries a 12 × 12 array of 144 microdomes sized for a 35 mm dish or a 6-well plate, and is trimmed down to, for example, 9 × 9 (81 domes) for an 8-well chamber or a 24-well plate (**Fig. 1c**); array size and pitch are set by the mold and can be redesigned for any format.

Since UV curing is integral to fabrication, we further verified that the Microdomes chips can be produced with UV equipment more commonly available across laboratories rather than the specific source used here. The chips were successfully fabricated using three alternative UV sources – a 3D-printing post-curing station, a UV Clave sterilising cabinet and even a nail-polish curing lamp – with the corresponding curing times reported for each **(Supplementary Fig. 1b – e)**. This allows other laboratories to fabricate their own devices using UV equipment that they already have. We further show that the Microdomes chip can be peeled off from its coverslip **(Supplementary Fig. 1f)**, allowing cultured samples to be recovered from the array for downstream molecular biology applications.

Prior to cell seeding, the Microdomes are surface-passivated with 0.5% Lipidure®-CM5206, a 2-(methacryloyoxy)ethyl phosphorylcholine (MPC)-based cell membrane-mimetic coating that prevents cell adhesion^13^. We later show that this passivation remains effective throughout long culture periods without apparent cytotoxicity, and that passivated devices can be stored for several months before use (refer to Methods for detail).

### A trap-like geometry shields specimens from shear and loss without restricting exchange

Any format intended for routine use must accommodate the liquid-handling operations that structure laboratory workflows — cells, medium, matrix and staining reagents delivered by pipetting, in parallel across many wells, which argues for a cavity left open at the top, with no lid or fluidic connection to the specimen. Such a chamber must then satisfy two opposing requirements: retaining a growing 3D specimen — an embryoid body, tumoroid or organoid — while allowing free exchange of soluble factors with the surrounding medium. Each Microdome resolves this through a 100 µm apical opening, wide enough for suspended cells to enter during seeding but too narrow for them to escape once they have aggregated into spheroids.

We first tested whether this opening limited molecular transport by tracking the entry of fluorescent dextran into Microdomes using confocal time-lapse imaging (**Fig. 2a**). Dextran of 10, 40 and 2000 kDa were selected to cover a broad molecular size range relevant to culture and downstream processing, including nutrients and metabolites, growth factors and cytokines, serum proteins, as well as antibodies used for immunostaining (i.e VEGF dimer, 45 kDa; albumin, 66 kDa; immunostaining antibodies IgG, 150 kDa; and matrix components fibronectin, 440 kDa; laminin-111, 900 kDa; high-molecular-weight hyaluronan, >1000 kDa). Following addition of dextran solution, fluorescence within Microdomes increased progressively for all molecular weights tested. Fitting the fluorescence accumulation curves using a one-phase association model yielded apparent equilibration half-times (t1/2) of 11.6 min for 10 kDa dextran, 18.4 min for 40 kDa dextran and 19.3 min for 2000 kDa dextran (**Fig. 2b**). A 200-fold increase in molecular mass therefore slowed equilibration by less than a factor of two, and even the largest tracer filled the cavity well within a single handling step, indicating that the apical opening is permissive across the full range of molecules used in 3D culture and immunostaining, not merely small solutes.

**Fig. 2.**
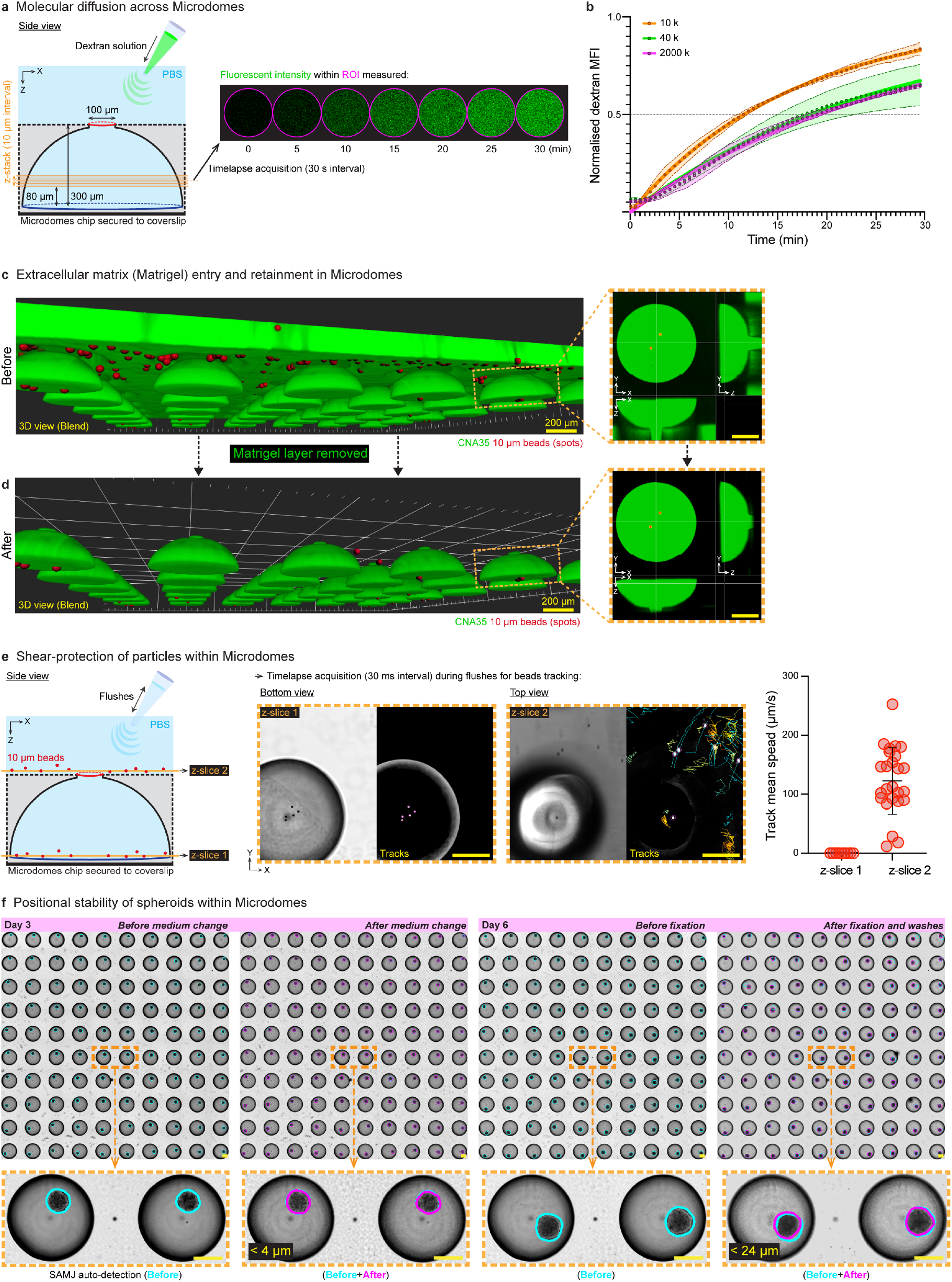
Fluidic characterisation of the Microdomes. **a.** Schematic of dextran diffusion assay. **b.** Quantification of dextran diffusion using 10 kDa, 40 kDa and 2000 kDa dextran. Mean fluorescence intensity (MFI) of dextran within ROI was normalised to the fitted baseline and plateau of a one-phase association fit. Circles, measured values; solid lines, one-phase association fits; shaded bands, mean ± SD (*n* = 5 z-positions within one dome per condition). The dashed line marks half-maximal accumulation. **c.** Images of CNA35-labelled Matrigel containing 10 µm fluorescent beads added to Microdomes. **d.** Images of Microdomes after removal of overlying Matrigel layer. **e.** Schematic of flushing assay for tracking bead movement. Tracking of bead movement during flushing was performed using TrackMate in FIJI, measured as the track mean speed (µm/s) of beads imaged inside (z-slice 1) and outside (z-slice 2) Microdomes (*n* = 25 (inside) and 9 (outside) tracks, mean ± SD). **f.** Assessment of spheroid positional stability within Microdomes during handling steps (*n* = 100 spheroids). Spheroid positions in the array were automatically detected using SAMJ in FIJI. Scale bar is 200 µm.

Extracellular matrices (ECM) supplementation provides the biochemical and structural cues that drive 3D growth and morphogenesis and is standard in most of 3D biology protocols. We therefore asked whether Microdomes could accommodate ECM. The choice of matrix is application-dependent, ranging from fibrin hydrogels for angiogenesis models^14^ to hyaluronic acid hydrogels for lung alveolar organoids^15^ and tissue-specific decellularized ECM for organoid cultures^16,17^. Here, we used Matrigel^18,19^ as a representative matrix to demonstrate the compatibility of Microdomes with matrix-based 3D culture. CNA35-labelled undiluted Matrigel containing 10 µm fluorescent beads was added onto Microdomes and allowed to enter the cavities on ice for 15 min. 3D reconstructions showed that the matrix entered the cavities together with the beads (**Fig. 2c**). Once the Matrigel had fully polymerised, the overlying matrix layer could be cleanly removed while preserving the Matrigel and embedded beads within the Microdomes (**Fig. 2d**). Thus, Microdomes can support local matrix retention while preserving direct access to the culture medium, avoiding the need for a continuous thick matrix layer above the array.

Handling-induced flow is a further constraint, particularly for the fragile spheroids and organoids produced by scaffold-free systems. Pipetting must be performed with care: the shear generated during fluid exchange can displace, disrupt or wash out specimens, and sublethal shear is itself a mechanical stimulus capable of altering cell phenotype^20,21^. We therefore tested whether the Microdome geometry shields particles within the cavity from transient flow during flushing (**Fig. 2e**, **left)**. Using 10 µm fluorescent beads as cell-sized tracers, time-lapse imaging qualitatively and quantitatively showed that beads outside Microdomes were rapidly displaced during flushing (123.6 ± 56.5 µm/s, *n* = 25 beads individually tracked), whereas beads inside Microdomes remained largely stationary (0.29 ± 0.07 µm/s, *n* = 9 beads individually tracked), corresponding to a 99.8% reduction in mean speed of beads (tracking and quantification performed using TrackMate ^22^; **Fig. 2e**, **right)**. This shear-protection property is important for the broad biological applicability of Microdomes: minimising handling-induced shear stress is particularly valuable during long-term 3D culture, where repeated medium changes are routinely required, and may also broaden the range of cell types that can be cultured, including shear-stress-sensitive populations such as suspension cells.

Finally, we examined whether this protection translated to positional stability of spheroids during routine experimental handling. To assess this, we imaged spheroid arrays in brightfield before and after two routine handling steps: a medium change at day 3, and fixation followed by multiple washing at day 6, using the same array (**Fig. 2f**). Spheroid positions were automatically segmented in batch using SAMJ Annotation in FIJI^23^ (Methods), and the before and after outlines were overlaid onto the corresponding brightfield images. After medium change, the outlines showed strong overlap, with mean centroid displacements below 4 µm (3.96 ± 2.41 µm, *n* = 100 spheroids) – under 3% of the mean spheroid diameter at that timepoint – while fixation and washing introduced only slight positional shifts (below 24 µm on average (23.6 ± 37.1 µm, *n* = 100 spheroids), which is approximately 15% of the mean spheroid diameter), as expected from the additional handling steps; the per-dome centroid displacements are shown in **Supplementary Fig. 2**, and spheroids nonetheless remained within their respective Microdomes. Together, these results show that Microdomes combine molecular accessibility, matrix compatibility and sample protection, supporting their use as stable microenvironments for scalable 3D culture and imaging workflows.

### Microdomes provide standardised 3D culture across cell types, laboratories and microscopes

To establish the Microdome as a versatile platform for 3D culture, we assessed its compatibility across a panel of nine cell lines spanning epithelial carcinomas, fibroblasts and neuroblastoma (**Fig. 3**, **Supplementary Fig. 3)**. As the Microdomes chips are fabricated using NOA, a mechanically robust and dimensionally stable polymer, passivated and sterilised devices could be prepared at one site and transported overseas to independent laboratories as ready-to-use devices. We show that the Microdomes supported spheroid formation across diverse handling protocols, cell types and imaging systems, and required no specialised equipment beyond the incubators and microscopes already in routine use, underscoring its practicality as an observation window for longitudinal monitoring of spheroids.

**Fig. 3.**
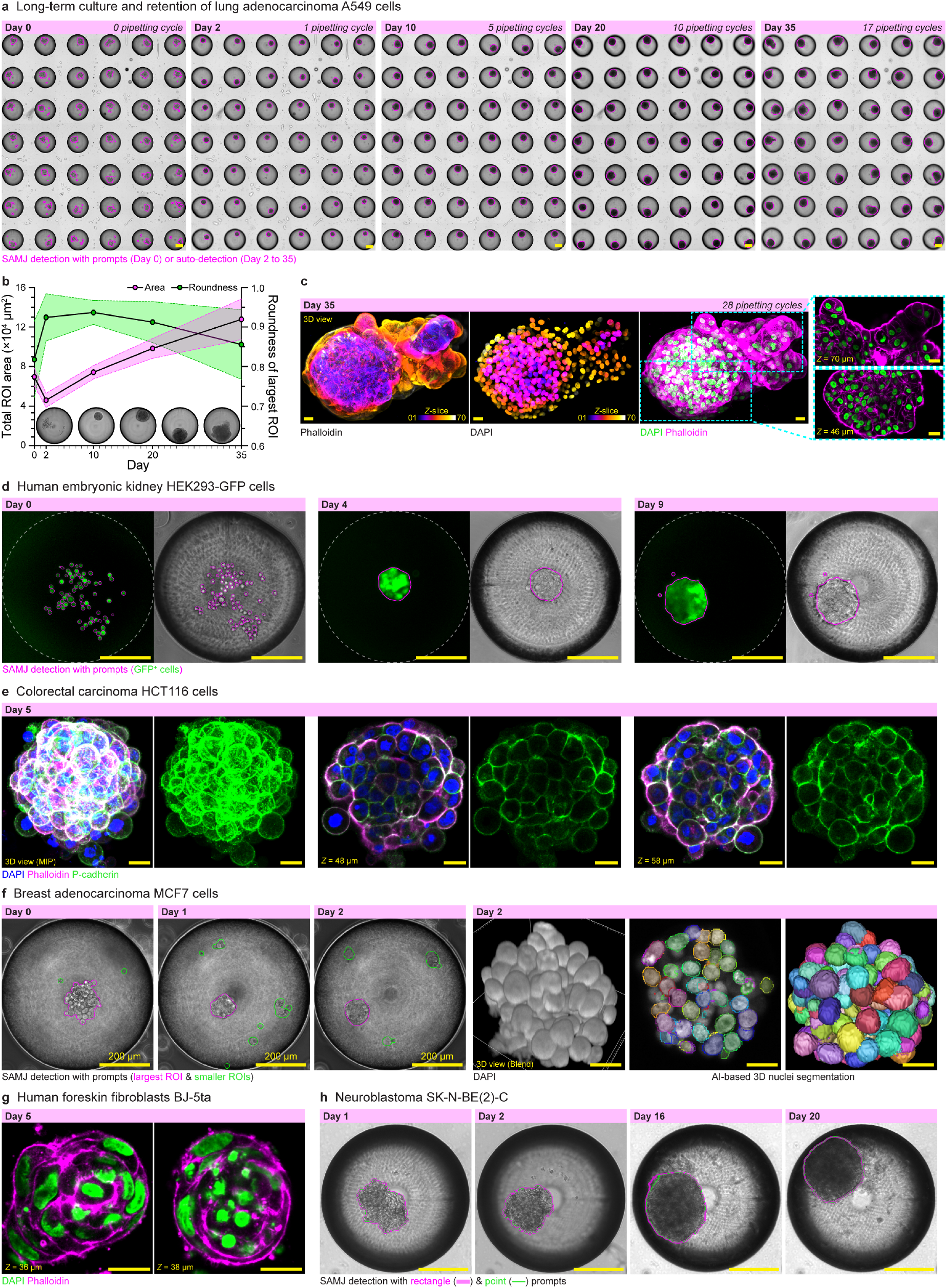
Microdomes support three-dimensional culture of diverse cell types. **a.** Long-term culture and retention of lung adenocarcinoma A549 spheroids tracked in a Microdome array from Day 0 to Day 35; the cumulative number of pipetting cycles is indicated at each time point. Spheroids were segmented with SAMJ (magenta), using manual prompts at Day 0 and automated detection from Day 2 onward, producing the ROIs used for quantification in b. **b.** Total ROI area (×10⁴ µm², magenta, left axis) and roundness of the largest ROI (green, right axis) across culture duration (*n* = 48 domes, mean ± SD; representative brightfield images of a single spheroid are shown below the plot). **c.** High-resolution 3D confocal imaging of a representative A549 spheroid at Day 35 stained for F-actin (phalloidin) and nuclei (DAPI). The phalloidin and DAPI channels are colour-coded by z-depth (z-slice 1–70); the merge shows DAPI (green) and phalloidin (magenta), with boxed regions displayed at z = 46 µm and z = 70 µm. **d.** HEK293-GFP spheroids at Day 0, 4 and 9, each from a different dome, imaged in fluorescence (GFP, green) and brightfield, with SAMJ detection of GFP⁺ cells overlaid (magenta); dashed lines indicate the dome boundary. **e.** Colorectal carcinoma HCT116 spheroid at Day 5 imaged by 3D confocal microscopy (maximum-intensity projection (MIP) and optical sections at z = 48 and 58 µm), immunostained for DAPI (blue), phalloidin (magenta) and P-cadherin (green). **f.** Breast adenocarcinoma MCF7 spheroids: brightfield time course of a single dome at Day 0, 1 and 2, with SAMJ detection of the largest ROI (magenta) and smaller ROIs (green), shown alongside a separate Day 2 spheroid imaged by 3D confocal microscopy of DAPI (3D view in blend mode) with AI-based 3D nuclei segmentation. **g.** Human foreskin fibroblast BJ-5ta spheroid at Day 5 (optical sections at z = 36 and 38 µm), stained for nuclei (DAPI, green) and F-actin (phalloidin, magenta). **h.** Neuroblastoma SK-N-BE(2)-C spheroids imaged in the same dome at Day 1, 2, 16 and 20 (brightfield), with SAMJ detection using rectangle (magenta) and point (green) prompts. Scale bars, 200 µm (a, d, h) and 20 µm (c, e, f, g), unless otherwise stated.

To assess compatibility with long-term culture, lung adenocarcinoma A549 cells were maintained in Microdomes for over one month before endpoint fixation and high-resolution 3D confocal imaging (**Fig. 3a–c**). Individual spheroids were reliably tracked from Day 0 to Day 35 (**Fig. 3a**) and were fully retained within their domes despite repeated pipetting during long-term maintenance, consistent with the shear protection demonstrated earlier (**Fig. 2e, f**). Both dispersed cells and subsequently compacted spheroids were segmented for morphological quantification using SAMJ in Fiji: at seeding (Day 0), prompts were placed manually for each dome, whereas from Day 2 onward the compacted spheroids were segmented using the batch workflow that automatically generated the prompts (**Fig. 3a–b**; Methods). Because every dome occupies a fixed position within the standardised arrangement, large numbers of spheroids could be readily followed in parallel across the full time course. This is demonstrated here for a representative array of 48 domes. The total segmented cell area decreased over Days 0 – 2 as cells aggregated and compacted and then increased steadily thereafter. During this initial compaction, spheroid roundness rose and was maintained through Day 20. It then declined gradually toward Day 35, coinciding with the emergence of small protrusions at the spheroid surface (**Fig. 3b**). Endpoint confocal imaging of DAPI- and phalloidin-stained spheroids resolved distinct cellular arrangements between the spheroid periphery and core, including columnar cells at the surface, and showed that the original rounded body remained densely packed whereas the later protrusions were more loosely organised (**Fig. 3c**), illustrating the subcellular detail accessible through the dome at the end of an extended time course.

Beyond A549 cells, the Microdomes supported spheroid formation by eight other cell lines: HEK293 immortalised embryonic kidney cells expressing GFP (HEK293-GFP), colorectal carcinoma HCT116, breast adenocarcinoma MCF7 and MDA-MB-231, BJ-5ta foreskin fibroblasts, and the neuroblastoma lines SK-N-BE(2)-C, SH-SY5Y and SK-N-AS (**Fig. 3d–h**, **Supplementary Fig. 3)**, across culture periods ranging from 2 to 20 days and using cell-type-specific seeding and medium conditions (Methods). For instance, the Microdomes readily accommodate seeding formats and conditions tailored to experimental constraints (e.g., Microdomes can be mounted on cultureware sized to suit the number of cells available; **Fig. 1c**) or cell-type-specific requirements (e.g., neuroblastoma suspensions were supplemented with methylcellulose to support aggregation).

As different cell types were monitored on different microscopes, we applied a single segmentation approach, SAMJ, to images from different platforms, confirming that spheroids could be reliably imaged and segmented for downstream analysis regardless of the acquisition system (**Fig. 3a, d, f, h**, **Supplementary Fig. 3)**. Endpoint staining and analysis were equally straightforward: HCT116 spheroids were immunostained for P-cadherin (**Fig. 3e**), and DAPI-stained MCF7 spheroids were segmented nucleus by nucleus in 3D with a StarDist convolutional neural network, as described previously¹² (**Fig. 3f**). Microdomes therefore act as a standardised, transportable observation window, one that travels between laboratories and microscopes, and delivers images that feed directly into existing analysis pipelines, without specialised handling or equipment.

### Microdomes hold 3D cultures in place to enable sequential co-culture within a single dome

Having established robust spheroid formation across diverse cell types (**Fig. 3**, **Supplementary Fig. 3)**, we next investigated whether Microdomes could support co-culture workflows, in which a second population must be introduced without perturbating the first (**Fig. 4**, **Supplementary Fig. 4)**.

**Fig. 4.**
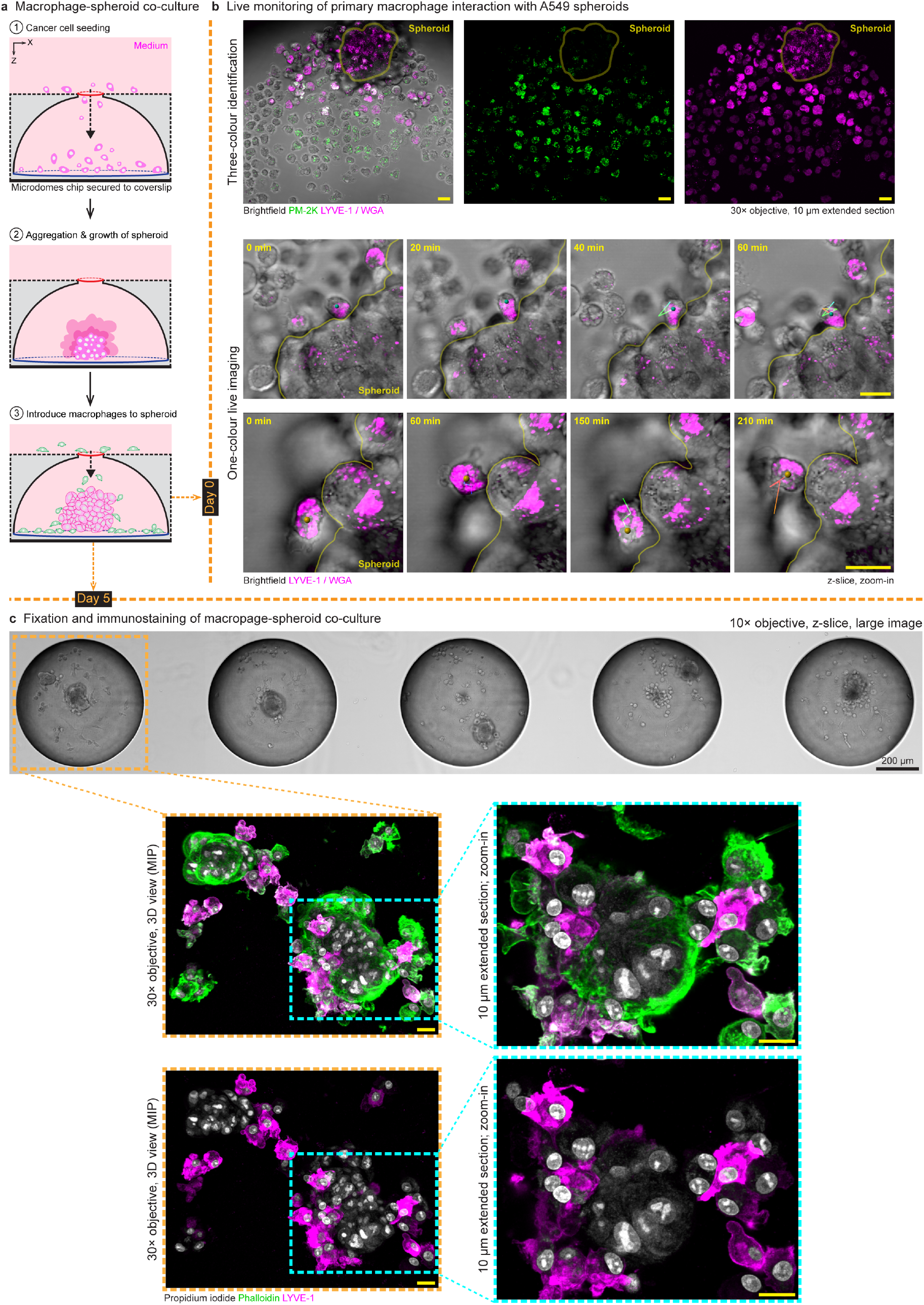
Microdomes support macrophage–spheroid co-culture. **a.** Schematic of the co-culture workflow. **b.** Live monitoring of primary macrophage interactions with A549 spheroids in Microdomes. A reference image acquired before the time-lapse distinguishes macrophages with three-colour identification (PM-2K, green; LYVE-1, magenta) from the WGA-labelled spheroid (magenta); live imaging was then performed in brightfield and the magenta channel only, with the spheroid boundary indicated by the yellow outline. Time-series images show two representative macrophage–spheroid interaction events; tracks indicate the movement of individual macrophages in contact with the spheroid. **c.** Immunostaining of fixed macrophage–spheroid co-culture in Microdomes. A section of a large-area brightfield image of the co-culture is shown, with the boxed Microdome displayed below by 3D confocal imaging. Scale bars, 20 µm unless otherwise stated.

We first confirmed compatibility with the simplest format, in which distinct cell types are seeded together and left to co-aggregate into a single spheroid, as shown for A549 cells co-seeded with BJ-5ta fibroblasts; the Microdomes optical access allowed both populations to be resolved individually within the aggregate **(Supplementary Fig. 4)**.

We focus below on the sequential format, in which one population is established before the other is introduced, using a macrophage-tumour spheroid co-culture to monitor direct interactions between the two cell types, whose relevance to tumour progression has been highlighted by earlier studies^24,25^. To demonstrate this, A549 cells were first seeded into Microdomes and allowed to form spheroids for 7 days. LYVE-1^+^ macrophages were separately differentiated from human donor-derived monocytes, purified by flow cytometry and introduced to the pre-formed spheroids (**Fig. 4a**). This sequential workflow requires the initial spheroids to remain stably positioned during subsequent cell addition, medium exchange and imaging, highlighting the importance of the ability of Microdomes to retain spheroids during handling, as demonstrated earlier (**Fig. 2e, f**).

To distinguish the two cell types, A549 spheroids and macrophages were labelled separately prior to co-culture. The A549 spheroid cell surface was labelled with membrane dye WGA, whereas macrophages were identified using PM-2K, a marker of human tissue macrophages, together with LYVE-1 marker to distinguish the LYVE-1^+^ macrophage subset (**Fig. 4b**, **top)**. Following macrophage entry into the Microdome cavities, live imaging was performed to monitor macrophage-spheroid interactions. Although the two cell types were morphologically distinct – macrophages as individual, motile cells and the spheroid as a compact aggregate with a smooth boundary – the separate fluorescence labelling allowed macrophages in contact with the spheroid to be unambiguously identified and tracked over time. Representative time-series images showed individual macrophages migrating along the spheroid surface while maintaining contact with the spheroid boundary (**Fig. 4b**, **bottom)**.

The co-culture was maintained for a further 5 days before fixation and immunostaining. Because each dome held one spheroid at a known position and its associated macrophages, every unit in the array shared the same geometry and the same accessible volume making them directly comparable. A single large-area brightfield scan captured the whole array at once (**Fig. 4c**, **top)**, and any individual unit could then be revisited at high resolution by 3D confocal microscopy (**Fig. 4c**, **bottom**). Zoomed-in extended-section images further resolved LYVE-1^+^ macrophage localisation around the spheroid and revealed protrusive macrophage morphologies at the spheroid interface (**Fig. 4c**, **bottom right)**.

Together, these results show that Microdomes accommodate both simultaneous and sequential co-culture, from live monitoring of heterotypic interactions to endpoint immunostaining — turning each dome into a self-contained, directly comparable unit for studying multicellular interactions.

### A fixed and optically transparent ‘address’ turns correlative microscopy into a routine operation

A key advantage of the Microdomes is that individual spheroids are held in a defined array and can be repeatedly relocated across imaging modalities, enabling the same 3D object to be tracked ‘longitudinally’ from initial cell aggregation through growth, fixation, staining, and high-resolution endpoint imaging.

To demonstrate this, A549 cells seeded into Microdomes were first followed by phase contrast live imaging over the first 24 h, on a widefield microscope (Nikon BioStation IM-Q), capturing the initial aggregation of seeded cells (**Fig. 5a**, **Supplementary Fig. 5a, Supplementary Movie 1)**. Over the following days, the aggregates developed into growing spheroids, and the array was captured as a brightfield large image on a spinning disk confocal microscope (BC43 Benchtop confocal microscope) at days 2, 4 and 6 (**Fig. 5b**, **Supplementary Fig. 5b)**. The regular array geometry allowed individual spheroids to be unambiguously relocated at each time point.

**Fig. 5.**
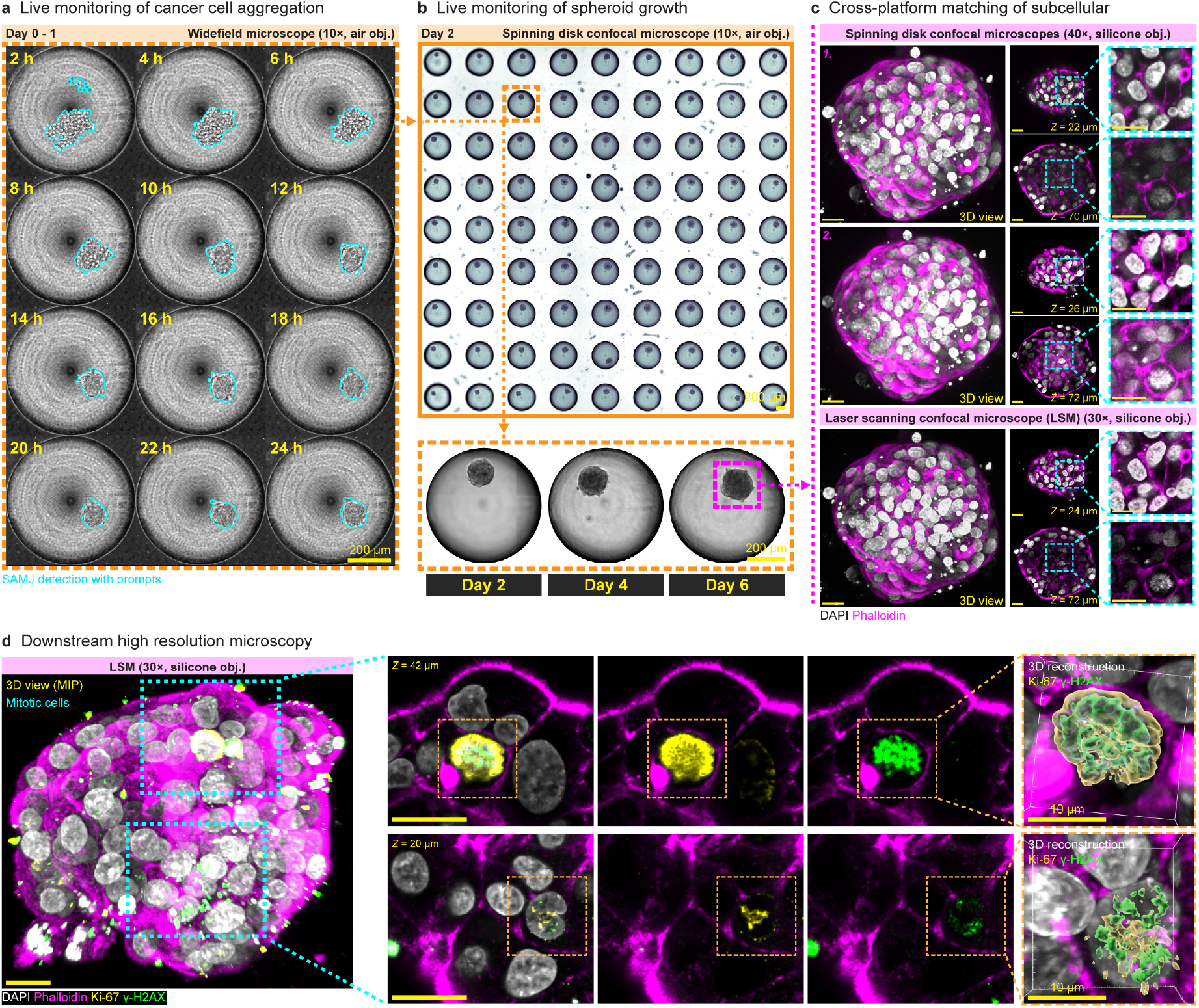
Microdomes support correlative microscopy of spheroids. **a.** Live widefield imaging of A549 cell aggregation 2 h-post seeding (using Nikon BioStation IM-Q, 10× air objective). Representative time-series images with cells and aggregates annotated by SAMJ in Fiji. **b.** Longitudinal large-area imaging of spheroid growth, on Day2, 4, and 6, within the Microdome array (using Andor BC43 benchtop spinning disk confocal, 10× air objective). **c.** Correlative imaging of longitudinally tracked spheroids across multiple confocal platforms: (1.) an Andor BC43 benchtop spinning disk confocal (40× silicone objective), (2.) a Yokogawa CSU-W1 spinning disk confocal on a Nikon Ti-E (40× silicone objective) and an Olympus FV3000 laser scanning confocal (30× silicone objective). Images of a representative spheroid (DAPI, white; phalloidin, magenta) are shown, with matched z-sections and zoomed regions highlighting corresponding subcellular features across platforms. **d.** Representative high-resolution laser scanning confocal (using Olympus FV3000, 30× silicone objective) imaging of spheroid (DAPI, white; phalloidin, magenta; Ki-67, yellow; γ-H2AX, green), with zoomed optical sections and three-dimensional reconstructions resolving nucleus-associated features of mitotic cells. Scale bar is 20 µm unless otherwise stated.

After 6 days of culture, the sample was fixed and stained for F-actin (phalloidin) and nuclei (DAPI). To test whether the device tolerates repeated downstream imaging and handling, the same sample was imaged on three confocal platforms – two spinning disk systems and one laser scanning microscope (refer to Methods for detail) – with the sample re-stained, washed and stored at 4 °C between each platform (**Fig. 5c**, **Supplementary Fig. 5c)**. Despite these repeated rounds of handling, the same spheroids were reliably relocated on all 3 microscopes.

Beyond relocating whole spheroids, subcellular landmarks within the spheroids were preserved and could be matched across platforms. Mitotic cells, for example, readily identifiable from their characteristic condensed chromatin, could be located in confocal z-stacks acquired on each microscope. As indicated by the z-positions annotated on each slice, matched structures were recovered at closely comparable depths within the spheroid across all 3 platforms, and the same was observed for six independently tracked spheroids **(Supplementary Fig. 5c)**. Consistent with the positional-stability measurements shown earlier (**Fig. 2f**), this indicates that the device protects spheroids from handling-associated shear, preserving both their position in the array and their internal architecture through repeated handling.

The workflow was also compatible with immunostaining for specific markers. As a proof of principle, spheroids were stained for proliferation marker Ki67 and DNA damage marker γH2AX, showing that marker-specific, high-resolution endpoint imaging can be appended to the correlative workflow within the device (**Fig. 5d**). High-resolution confocal z-stacks resolved the organisation of Ki67 and γH2AX signal within individual mitotic cells, and 3D segmentation and reconstruction of these signals confirmed that subcellular structures within spheroids can be clearly resolved.

Together, these results establish the Microdomes as a platform for correlative microscopy, linking live observation of spheroid/embryonic bodies formation, array-based relocation, repeated imaging across multiple microscope platforms, and high-resolution immunofluorescence of defined subcellular structures. The ability to recover not only the same spheroids but the same subcellular landmarks after repeated handling underscores the utility of the device for longitudinal, multimodal, endpoint-correlative imaging.

## Discussion

Microdomes hold together, in a single device, properties that existing 3D culture formats achieve only in part (**Table 1**): platforms that fix specimens at addressable positions leave them exposed to flow or behind a millimetre of plastic, while those that protect them do so through dedicated ports rather than through the geometry itself. Shielding specimens from flow-induced shear keeps them in place through direct top-down pipetting, yet the open apex leaves them accessible to soluble macromolecules, imaging reagents and, where needed, physical retrieval — all while remaining on top of and near a conventional 170 µm coverslip. Because protection and accessibility no longer compete, a culture can stay in the same device from seeding through growth, perturbation, fixation and imaging, and be interrogated at every scale along the way, from array-wide time courses to the subcellular organisation of one specimen.

**Table 1.**
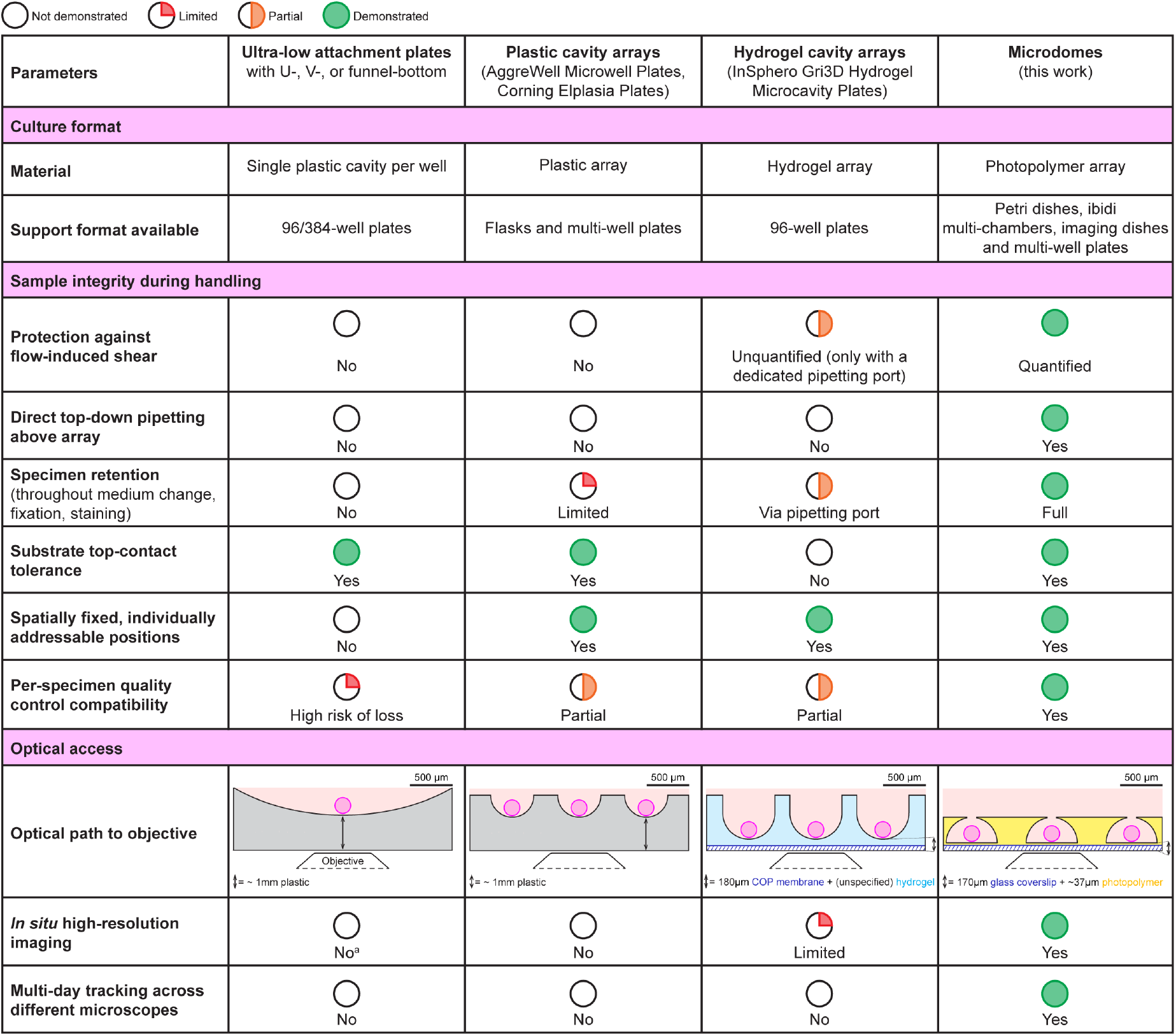
Microdomes in the context of existing (micro)well technologies. Entries for the Microdomes are supported by data in this study; entries for the other systems are based on manufacturer documentation and published literature^3,4^. All schematics are drawn to the same scale; the objective is a symbol and is not to scale. ^a^ Thick-bottom ultra-low attachment plates (∼1 mm) are considered here. Plates with a thin optical bottom, such as the Akura™ 96/384 Spheroid Microplate (125 µm), permit *in situ* high-resolution imaging but share the other limitations listed.

| **Parameters** | **Ultra-low attachment plates**<br>with U-, V-, or funnel-bottom | **Plastic cavity arrays**<br>(AggreWell Microwell Plates, Corning Elplasia Plates) | **Hydrogel cavity arrays**<br>(InSphero Gri3D Hydrogel Microcavity Plates) | **Microdomes**<br>(this work) |
|---|---|---|---|---|
| **Culture format** |  |  |  |  |
| **Material** | Single plastic cavity per well | Plastic array | Hydrogel array | Photopolymer array |
| **Support format available** | 96/384-well plates | Flasks and multi-well plates | 96-well plates | Petri dishes, ibidi multi-chambers, imaging dishes and multi-well plates |
| **Sample integrity during handling** |  |  |  |  |
| **Protection against flow-induced shear** | No | No | Unquantified (only with a dedicated pipetting port) | Quantified |
| **Direct top-down pipetting above array** | No | No | No | Yes |
| **Specimen retention**<br>(throughout medium change, fixation, staining) | No | Limited | Via pipetting port | Full |
| **Substrate top-contact tolerance** | Yes | Yes | No | Yes |
| **Spatially fixed, individually addressable positions** | No | Yes | Yes | Yes |
| **Per-specimen quality control compatibility** | High risk of loss | Partial | Partial | Yes |
| **Optical access** |  |  |  |  |
| **Optical path to objective** | 500 µm<br>Objective<br>↕= ~ 1mm plastic | 500 µm<br>↕= ~ 1mm plastic | 500 µm<br>↕= 180µm COP membrane + (unspecified) hydrogel | 500 µm<br>↕= 170µm glass coverslip + ~37µm photopolymer |
| ***In situ* high-resolution imaging** | No<sup>a</sup> | No | Limited | Yes |
| **Multi-day tracking across different microscopes** | No | No | No | Yes |

More than 100 3D biological objects can be maintained on a single chip and quantitatively tracked over extended periods using automated segmentation and tracking, while the chip’s thin, transparent construction enables high-resolution 3D confocal imaging and precise relocation of the same microstructures across microscopy platforms. Because each specimen keeps the same address for the lifetime of the culture, every dome can be followed individually from seeding to endpoint, so imaging becomes per-specimen quality control rather than a snapshot of a population: batch homogeneity, growth trajectories and outliers can be documented for every unit produced.

Combined with the shear shielding that preserves specimen integrity and minimises unwanted phenotypic perturbation during repeated handling, this addresses two practical requirements of organoid production at scale, specimen survival and specimen-level traceability, that are not met by formats in which samples move, merge or are lost between time points. Importantly, Microdomes are designed for broad compatibility with standard laboratory workflows, from fabrication and cell culture to imaging and analysis. The chips can be produced using accessible fabrication methods, integrated into common culture formats, used with diverse cell types, and imaged across conventional microscopy platforms. Together, these features position the device as a versatile and readily adoptable interface between routine 3D culture and multimodal analysis.

The current implementation nevertheless has limitations that arise primarily from its fabrication and assembly. Although individual chips can be fitted into several standard culture vessels, individual placement become increasingly difficult as the culture format is miniaturised. Automation of production to accommodate 96-well plates format, massively used in drug screening, will be helpful to tackle this limitation.

Beyond the configuration demonstrated here, the Microdomes concept provides a modular framework that could be adapted to a broader range of 3D culture applications. NOA was selected because it combines optical clarity with mechanical robustness and supports prolonged culture and repeated handling. Preliminary testing also suggests that assembled devices can tolerate freeze-thaw procedures of cryopreservation, raising the possibility of preserving spatially indexed 3D culture libraries for biobanking. More broadly, the fabrication strategy is not restricted to NOA or glass coverslip. Dome arrays could potentially be produced from alternative materials, including PEG hydrogels or decellularized ECM, to provide defined biochemical or mechanical tuneable cues.

Likewise, attachment of chip to porous membranes or plastic culture substrates could extend the platform to barrier models, compartmentalised co-cultures and standard disposable cultureware. These adaptations will require systematic validation, particularly with respect to material compatibility and lifetime. The same geometry should also open the specimens to imaging modalities not tested here — light-field, oblique light-sheet and two-photon microscopy, and potentially electron microscopy — further extending the range of volumetric readouts obtainable *in situ*. Taken together, these features make Microdomes an essential platform for high-throughput organoid culture – transforming routine organoid and spheroid assays into fully trackable, imageable resources for translational research, all within a single device capable of multiscale assessment from whole arrays down to the subcellular level.

## Acknowledgments

A.B. acknowledges funding support from the Mechanobiology Institute (MBI) seed grant. G.G. acknowledges MBI for the support. M.L.G. and E.P. acknowledge funding support from La Ligue contre le cancer (Equipes labellisées 2025). K.P. acknowledges funding support from the NSW Space Research Network (with P.D.) and the US Department of Airforce, AOARD (FA2386-23-1-4045). S.E.G. acknowledges internal funding from the Center for CardioVascular and Nutrition Research (C2VN).

V.A. was supported by a seed fund from MBI, NUS and the French government through the France 2030 investment plan managed by the National Research Agency (ANR), as part of the Initiative of Excellence Université Côte d’Azur under reference number ANR-15-IDEX-01 and through Chair Professor Junior by INSERM. V.A. and N.L.S.J are currently funded by a grant “Leader in Oncology” from the Fondation ’ARC pour la recherche sur le cancer’. Y.T. acknowledges funding support from National Research Foundation of Singapore under its mid-sized grant (NRF-MSG-2023-0001).

We thank the MBI Microfabrication and Microscopy Cores for technical support with device production and imaging, and Femtoprint SA (Agno, Switzerland) for the fabrication of the glass primary mold. We are grateful to J. Young and R. Arulselvam from MBI for the gift of MDA-MB-231 cells, and to the healthy blood donors and the NUS blood-donation programme for the apheresis cones used to derive primary human macrophages. Imaging at UNSW was performed at the Katharina Gaus Light Microscopy Facility (KGLMF), Mark Wainwright Analytical Centre, UNSW Sydney. We would like to thank Dr. Alison Ferguson from the 3D culture facility of KGLMF, UNSW for her contribution to the testing of the microdevices and for her feedback during the revision of the manuscript and figures.

## Authors contribution

A.B. and G.G. conceived the study and designed the Microdomes. G.G. developed the fabrication process and wrote the microfabrication sections. M.S.C., S.B.M.R., E.K. and A.B. fabricated, passivated and characterised the devices. M.S.C., J.Liu and J.C.F.L. performed the dextran diffusion, Matrigel incorporation, flushing and positional-stability experiments. M.S.C., M.G. and A.B. performed spheroid culture and imaging for the cell-line panel; M.G. performed the fibroblast co-culture experiments; S.E.G. performed the HEK293-GFP experiments at AMU; M.L.G and E.P. performed the neuroblastoma cultures at CRCM; P.D. and K.P. performed the MCF7 cultures at UNSW. N.L.S.J., J.L.T.C. and V.A. produced and characterised the primary human macrophages and designed the co-culture experiments. H.T.O. and A.B. developed the segmentation, 3D analysis and tracking pipelines. Y.T. and M.G. contributed to the writing of the manuscript. A.B. supervised the study and wrote the manuscript with input from all authors. All authors read and approved the final manuscript.

## Data availability

Source data underlying the figures are provided with this paper. Due to storage space restrictions, the complete imaging datasets (raw and segmented 3D images, and the associated measurement tables) are available from M.S.C. and A.B. upon reasonable request; all requests will be answered within 2 weeks.

## Methods

### Fabrication methods for the Microdomes chips

First, the glass primary mold (Femtoprint SA, Agno, Switzerland) was replicated in polydimethylsiloxane (PDMS). A 10:1 ratio of PDMS prepolymer base to curing agent (Sylgard 184, Dow Corning) was mixed and degassed under vacuum (for 10 min at 10 mbar). The mixture was then poured onto the glass mold and degassed again (for 10 min at 10 mbar) to ensure complete filling of the microstructures. Following thermal curing (for 1 h at 80 °C), the PDMS replica mold was peeled off and cut into ∼1.5 × 1.5 cm² pieces, which served as reusable molds for fabricating Microdomes chips.

Next, a piece of 1.5 × 1.5 cm^2^ PDMS mold was inverted onto a cleaned glass slide or a flat PDMS substrate, and a small volume of UV-curable optical resin (NOA-73, Norland Optics) was dispensed at one edge of the mold. By capillary action, the liquid NOA-73 filled the spaces between the PDMS mold and the flat substrate. NOA was cured by UV exposure (UV LED KUB2, Kloe France, 365 nm at 100 mW/cm^2^ for 1 min). Subsequently, the PDMS mold was peeled off, and the sides of NOA-Microdomes chip (inverted) can be trimmed. Separately, glass coverslip of final culture vessel for the chip was coated with a thin layer of NOA-73, which was partially cured by UV exposure (at 40 mW/cm^2^ for 4 s), and served as adhesive for the chip. The trimmed NOA-Microdomes chip was then flipped onto the coated coverslip, and the entire set-up was exposed to UV (at 100 mW/cm^2^ for 1 min) as the final curing step.

### Long-term passivation of Microdomes chips

To prevent cell adhesion (validated up to 35 days) and allow spheroid formation, Lipidure (CM5206, NOF America) was used to passivate the Microdomes chip secured to glass coverslip. The device was first plasma treated (for 3 min) before 0.5% Lipidure (w/v) in 100% ethanol was added to sufficiently cover the chip. The device was then degassed (for 5-10 min at 10 mbar). Once the chip has been completely degassed, excess Lipidure solution was removed and completely evaporated (for at least 20 min at 37 °C), which gives rise to a cell membrane-mimetic coating. The passivated device was heat sterilised (for 20 min at 100 °C). From here on, the device is handled under sterile conditions.

Sterile PBS was added to the device, degassed (for 10 min at 10 mbar) and then filled with cell culture medium before cell seeding.

### Estimation of NOA-73 bonding-layer thickness

Coverslips were coated with fibrinogen-Cy5 (for 2 h at room temperature) to mark the coverslip surface, and bonded Microdomes (fabrication detailed above) were filled with calcein-containing solution (40 µg/mL in PBS). Confocal z-stacks were acquired using Andor BC43 benchtop confocal microscope (10× air objective, NA 0.45, 488 and 647 nm illumination, 2 µm z-step). For each dome, fluorescence intensity profiles were extracted from a circular ROI (37.1 µm diameter) near the dome centre using Plot Z-axis Profile in Fiji. Then, the fibrinogen-Cy5 profile was fitted with a Gaussian function to define the coverslip position, while the rising calcein profile was fitted with a sigmoid to define the Microdome base (Prism 11, GraphPad). The axial separation was corrected for refractive-index-induced scaling using the paraxial approximation (cured NOA-73, n = 1.56). For visualisation, z-positions within NOA-73 layer and the aqueous solution were corrected using scaling factors of 1.56 and 1.33, respectively, and fluorescence intensities were normalised.

### Dextran diffusion assay

Microdomes chip was secured to the centre of each well in a 6-well plate containing 5 mL of PBS. Fluorescein-dextran (10, 40 or 2000 kDa; D1821, D1844 or D7137, respectively, Thermo Fisher Scientific) was prepared at 100 µM, and 50 µL was pipetted into the PBS from the edge of the well. Time-lapse acquisition was started immediately upon dextran addition. For each dextran molecular weight, a z-stack of a central microdome was acquired at 80, 90, 100, 110 and 120 µm above the upper surface of the coverslip, every 30 s for 30 min, on a Zeiss laser-scanning confocal microscope (LSM) 980 (20× air objective, NA 0.8, 488 nm illumination, 10 µm z-step). Mean fluorescence intensity (MFI) was measured in Fiji from a circular ROI (138.11 µm diameter) positioned consistently in XY across all time points and z-positions. For each z-position, MFI measured before dextran addition was subtracted from all subsequent time points to obtain background-corrected fluorescence, which was fitted with a one-phase association model with Y0 unconstrained (Prism 11, GraphPad) to obtain the half-times. For visualisation, background-corrected fluorescence values were transformed to fractional accumulation as (Y − Y0)/(plateau − Y0) using its own fitted Y0 and plateau values, and the same model was re-fitted to the transformed data.

### Matrigel incorporation into Microdomes

Microdomes chip was secured to each well in a µ-Slide 8-well chamber (Ibidi) containing 200 µL of PBS. First, PBS was aspirated and 200 µL of a mixture of Matrigel (354277, BD Biosciences), EGFP-CNA35 (0.8 µM; a collagen-binding probe) and 10 µm red fluorescent FluoSpheres polystyrene microspheres (F8834, Thermo Fisher Scientific; 1.08 × 10⁵ particles) was added to each well. After 15 min on ice to allow the mixture to enter the domes, the Matrigel mixture was solidified at 37 °C overnight. To remove the overlying Matrigel layer, samples were washed three times with ice-cold PBS. Samples were imaged on an Andor BC43 benchtop confocal microscope (10× air objective, NA 0.45, 488 nm illumination, 5 µm z-step) before and after Matrigel layer removal.

### Flushing assay for bead tracking

Microdomes chip was secured to the centre of a µ-Dish 35 mm (Ibidi) containing 2 mL of PBS. 10 µm red fluorescent FluoSpheres polystyrene microspheres (1.08 × 10⁵ particles) were added to the PBS and resuspended to ensure homogeneous distribution. After 5 min, allowing enough beads to settle into the domes, brightfield time-lapse imaging was performed at two focal planes (one focused on beads near the base of Microdome cavities and one focused on beads outside the cavities, close to the upper surface of the Microdomes), during repeated manual flushing of the solution by pipette. Imaging was performed on Andor BC43 benchtop confocal microscope (10× air objective, NA 0.45, at 30 ms intervals for 15 s).

Bead tracking was performed in FIJI using TrackMate. Beads were detected using the LoG detector with settings adjusted for each focal plane to optimise bead detection. Track mean speed was extracted from TrackMate and only tracks with durations ≥6 s were retained for plotting.

### Cell lines and their maintenance

A549 lung adenocarcinoma cells (ATCC CCL-185), HCT116 colorectal cancer cells (Sigma-Aldrich 91091005), MDA-MB-231 breast adenocarcinoma cells (ATCC HTB-26), BJ-5ta foreskin fibroblast (ATCC CRL-4001), were cultured in complete DMEM [Dulbecco’s Modified Eagle Medium high glucose; Gibco 11965092 supplemented with 10% fetal bovine serum (FBS; Invitrogen 10082147) and 1% penicillin–streptomycin (Invitrogen 15070063)].

MCF7 breast adenocarcinoma cells (ATCC HTB-22) were cultured in complete EMEM [Eagle’s Minimum Essential Medium; supplemented with 10% FBS, 1% L-glutamine, 1% penicillin– streptomycin and 0.01 mg/ml human recombinant insulin (Sigma-Aldrich)].

HEK293 human embryonic kidney cells were a kind gift from Dr. Franck Peretti. Cells were transfected with a pCMV-C-GFPSpark plasmid (Sino Biological) using DreamFect Gold transfection reagent (OZ Biosciences) according to the manufacturer’s protocol, with 1 µg of plasmid DNA per well.

Transfected cells were maintained in DMEM (Gibco 11965092) supplemented with 10% FBS (Gibco 10270106), 0.5% penicillin-streptomycin (Gibco 15140122), 1% non-essential amino acids (Gibco 11140050) and hygromycin B (100 µg/ml; Gibco 10687010). All cell cultures were monthly screened to ensure the absence of mycoplasma contamination (MycoAlert® Assay LONZA and #LT07-518).

SH-SY5Y (RRID: CVCL_0019), SK-N-BE(2)-C (RRID: CVCL_0529) and SK-N-AS (RRID: CVCL_1700) neuroblastoma cell lines were obtained from European Collection of Cell Cultures. Upon receipt, cell master stocks were prepared and cells for experiments were passaged for less than 3 months.

Neuroblastoma cell lines were cultured in DMEM (Life Technologies 41965-039) supplemented with 10% FBS (Sigma-Aldrich GE F0392), 1% sodium pyruvate (Thermo-Fisher 11360070) and 1% penicillin/streptomycin (Thermo-Fisher 15070063). All cell cultures were monthly screened to ensure the absence of mycoplasma contamination (Eurofins Genomics or kit MycoAlert® Assay LONZA #LT07-710 and #LT07-518).

All cell cultures were maintained at 37 °C and 5% CO₂.

### Spheroid formation by different cell lines

Prior to seeding of A549, HCT116, MDA-MB-231 or BJ-5ta cells, chips were washed twice with complete DMEM. Each cell type was introduced at a range of 2-10 × 10^6^ cells/mL and incubated for 15-30 min (at 37 °C and 5% CO₂) to allow cells to enter the domes, with gentle swirling to resuspend settled cells at ∼ 10 min intervals. Excess cells were removed by 2 washes with PBS.

Prior to seeding of neuroblastoma cell lines, chips were washed thrice with PBS supplemented with antibiotic-antimycotic (Gibco 15240-062) followed by 3 washes with culture medium. Each cell type was introduced at 8 × 10^6^ cells/mL with 5% methylcellulose (Sigma M0512-100G) and incubated for ∼ 20 min (at 37 °C and 5% CO₂) to allow cells to enter the domes, with several rounds of pipetting to maintain cells in suspension. Excess cells were removed by 3 washes with culture medium.

Prior to seeding of HEK293 cells, chips were washed thrice with PBS supplemented with antibiotic-antimycotic (Gibco 15240-062) followed by 3 washes with culture medium. Each cell type was introduced at 5 × 10^6^ cells/mL and incubated for 5 min (on orbital shaker at room temperature) followed by a further 10 min incubation (at 37 °C and 5% CO₂) to allow sufficient cell entry. Excess cells were removed by 2 washes with culture medium.

Prior to seeding of MCF7 cells, chips were washed twice with culture medium. MCF7 cells were introduced at a range of ∼ 2.2 × 10^6^ cells/mL and incubated for 20 min (at 37 °C and 5% CO₂) to allow cells to enter the domes. Excess cells were removed by a replacement with culture medium.

Across all cell types tested, occupancy was typically 50-90 cells per dome. For all spheroid cultures, fresh medium was added every 2-3 days.

### Assessment of spheroid positional stability within Microdomes during handling

After A549 spheroids were cultured for 3 days, large-area tile-scan brightfield images (large images) of the array were acquired before and after the medium change. After a further 3 days of culture, spheroids were washed once with PBS, fixed in 4% formaldehyde (Thermo Fisher Scientific 28908) for 15 min and washed 3 times with PBS. Large images were acquired before and after this fixation and washing sequence, followed by segmentation to assess spheroid displacement (detailed below).

### Primary human macrophages

Peripheral blood mononuclear cells (PBMCs) were isolated from healthy-donor apheresis cones under protocols approved by the National University of Singapore Institutional Review Board (NUS-IRB-2022-345), with informed consent from all participants.

PBMCs were isolated using SepMate™-50 tubes and Lymphoprep™ (STEMCELL Technologies) according to the manufacturer’s instructions, and residual erythrocytes were removed by ACK lysis.

Classical monocytes were negatively enriched using the EasySep™ Human Monocyte Isolation Kit (STEMCELL Technologies), and purity (CD14^+^CD16^-^) was confirmed by flow cytometry. Monocytes were seeded at 1.2 × 10^7^ cells per T75 flask in X-VIVO 15 medium (Lonza) supplemented with M-CSF (40 ng/mL). After 24 h, cells were differentiated in MDI polarisation medium [M-CSF (100 ng/mL), dexamethasone (40 ng/mL) and IL-4 (10 ng/mL)], with medium replenished every 3 days^26^. On day 7, cells were harvested and stained with mouse anti-human CD45-BUV395 (BD Biosciences 563792), biotinylated mouse anti-human PM-2K (OriGene BM4037B) and goat anti-human LYVE-1 (R&D Systems AF2089), followed by Streptavidin–Pacific Blue (Thermo Fisher Scientific S11222) and donkey anti-goat Alexa Fluor 647 (Jackson ImmunoResearch) to detect PM-2K and LYVE-1, respectively. Dead cells were excluded using DAPI (KPL Scientific). LYVE-1^+^ macrophages were isolated as CD45^+^PM-2K^+^LYVE-1^+^ cells by fluorescence-activated cell sorting (FACSAria Fusion, BD Biosciences).

### LYVE-1⁺ macrophage-A549 spheroid co-culture

Microdomes chip was secured to each well in a µ-Slide 8-well chamber containing A549 spheroids cultured for 7 days. Prior to LYVE-1⁺ macrophage seeding, A549 spheroids were washed once with Hanks’ Balanced Salt Solution (Thermo Fisher Scientific 14025092) and stained with Alexa Fluor 647-conjugated WGA (wheat germ agglutinin; 5 µg/mL in HBSS) for 15 min at 37 °C. Spheroids were washed twice with PBS before replacement with complete DMEM. Subsequently, LYVE-1^+^ macrophages were introduced at 3 × 10^4^ cells per well in high-glucose DMEM supplemented with 10% human AB serum (Gibco), 1% penicillin–streptomycin (Gibco) and MDI polarisation factors. The co-culture was incubated for ∼ 20 min (at 37 °C and 5% CO₂) to allow macrophages to enter domes. Live imaging was performed on Olympus FV3000 (UPlanSAPO 30×S/1.05 silicone-immersion objective, 0.8-mm working distance, 2 µm z-step, 10 min intervals for 24 h) equipped with a stage-top incubator maintained at 37 °C and 5% CO2. The co-cultures were maintained for 5 days before fixation and immunostaining (detailed below).

### LYVE-1⁺ macrophage tracking in co-culture with A549 spheroids

Individual LYVE-1^+^ macrophages in contact with the spheroid were tracked manually in Imaris 10.2.0 (Oxford Instruments). Macrophages were identified from the pre-time-lapse reference image and their positions followed across the time-lapse using spot tracking. A549 spheroids in representative time series images were manually outlined using line tool in Fiji.

### On-chip immunofluorescent staining

For immunofluorescent staining, samples were washed once with PBS before fixing with 4% formaldehyde (Thermo Fisher Scientific 28908) for 15-20 min at room temperature. Fixed samples can be stored at 4 °C after PBS washes.

For LYVE-1, Ki-67 and γ-H2AX staining, samples were blocked and permeabilized in 5% BSA (Thermo Fisher Scientific 37525) with 0.2-0.3% Triton X-100 (Sigma-Aldrich T9284) for 30-60 min at room temperature. Primary antibodies, goat anti-human LYVE-1 (at 1:150; R&D Systems AF2089), rabbit anti-human Ki-67 (at 1:1,000; Abcam, ab15580), mouse anti-human phospho-Histone H2A.X (at 1:200; Cell Signalling 80312), were diluted in the blocking buffer and incubated at 4 °C overnight. For P- and N-cadherin staining, samples were permeabilized with 0.2% Triton X-100 for 30 min followed by blocking with 2% BSA in PBST for 1 h at room temperature. Primary antibodies, mouse anti-human P-cadherin (at 1:200; Thermo Fisher Scientific 324000) and rabbit anti-human N-cadherin (at 1:200; Cell Signalling 13116), were diluted in the blocking buffer and incubated at room temperature overnight.

Secondary antibodies, donkey anti-goat Alexa Fluor 647 (Jackson ImmunoResearch), donkey anti-rabbit Alexa Fluor 647 (Thermo Fisher Scientific A31573), donkey anti-mouse Alexa Fluor 568 (Thermo Fisher Scientific A10037), goat anti-mouse Alexa Fluor 488 (Thermo Fisher Scientific A11001) and goat anti-rabbit Alexa Fluor 488 (Thermo Fisher Scientific A11008), diluted at 1:200 and phalloidin (Thermo Fisher Scientific A12379 or A34055 or A22287) diluted at 1:100-1:400 in respective blocking buffer were added to samples and incubated at room temperature for 2 h, for LYVE-1, Ki-67 and γ-H2AX staining, or overnight, for P- and N-cadherin staining. Nuclei were counter-stained with DAPI (1:1,000; Thermo Fisher Scientific 62248) or Hoechst 33342 (1:1,000; Thermo Fisher Scientific H3570) or propidium iodide (1:1,000; Sigma-Aldrich P4864) at 1: 1,000 dilution in PBS for 20 min at room temperature. After each staining steps, samples were washed 3 times with PBS.

DAPI staining in MCF7 spheroids was performed with permeabilization using 1% Triton X-100, followed by blocking with 3% FBS, before incubation with DAPI (1:400) in PBS, with each step performed at 4 °C for 24 h. Samples were washed with PBS and incubated with a clearing solution (9 g of N,N,N,N-Tetrakis(2-Hydroxyproyl)ethylenediamine solution, 22 g of urea powder, 44 g of sucrose powder, 0.1 g of Triton X-100 and 24.9 g of deionized water) for 1 h prior to imaging.

### Image acquisition

Unless otherwise mentioned, spheroids from different cell types were acquired on different microscopes available at different research institutions.

HEK293 spheroids were imaged on EVOS M3000 fluorescence microscope (10× objective). MCF7 spheroids were imaged on Olympus IX71 (10× objective) and Leica Stellaris 8 (20× air objective, 1 µm z-step). Neuroblastoma spheroids were imaged on Leica DMIL LED® (4×/0.1 or 10×/0.1 objective). All other confocal z-stacks were acquired on Olympus FV3000 (UPlanSAPO 30×S/1.05 silicone-immersion objective, 0.8-mm working distance, 2 µm z-step). All 2D large images were acquired on Andor BC43 benchtop confocal microscope (10×/0.45 air objective).

### Correlative microscopy

Longitudinal tracking of same spheroids was performed across different microscope systems. 2 h after A549 cells were seeded, samples were monitored live on widefield Nikon BioStation IM-Q (10× air objective, NA 0.5, red LED illumination, 5 µm z-step spanning 65 µm, 1h interval for 24 h). Large images of spheroid array, including the tracked spheroids, were acquired on Day 2, 4, and 6. On day 6, spheroids were fixed and stained with DAPI and phalloidin (detailed above) for correlative imaging. Spheroids were stored at 4 °C for 1-12 days between acquisition on different microscope systems, with repeated staining and washing steps performed before each imaging. Confocal z-stacks (with 2 µm z-step) of same spheroids were imaged on Andor BC43 benchtop spinning disk confocal (40×/1.25 NA silicone-oil immersion objective), Yokogawa CSU-W1 spinning disk confocal on a Nikon Ti-E (40×/1.25 NA silicone-oil immersion objective) and an Olympus FV3000 LSM (UPlanSAPO 30×S/1.05 silicone-immersion objective).

### Spheroid segmentation with SAMJ

Prior to segmentation, related large images from the same experiment were aligned in Fiji using Linear Stack Alignment with SIFT (Similarity transformation model, using default settings). Spheroids in large images were segmented using a custom batch analysis macro for automatic segmentation with the Segment Anything Model (SAM) implemented as the SAMJ Annotator plugin in Fiji. Briefly, image pre-processing steps in the custom batch analysis macro produce one seed point per spheroid which are passed as prompts for SAMJ, using the EfficientSAM model, to generate spheroid outline ROI. Other than large images containing aggregated cells and spheroids, segmentation with SAMJ Annotator (EfficientSAM model) was performed with manually provided rectangle or point prompts.

### Spheroid tracking

ROI measurements were analysed using a custom Python pipeline. Each dome was assigned a deterministic identifier according to its position in the array, ordered top-to-bottom and left-to-right. ROIs were mapped to dome identifiers by their spatial positions and verified against the corresponding reference screenshots provided. In cases where multiple ROIs occupied one dome (before spheroid formation), all were retained under the same identifier. The pipeline compiles ROI-level dataset containing the source image, ROI label, dome identifier and available numeric Fiji measurements, from which dome-level summaries were calculated.

### Spheroid Displacement Analysis

To assess spheroid positional stability, paired before- and after-handling images were analysed using centroid coordinates from the Fiji ROI measurements. ROIs from the two images were assigned to the same dome identifiers, and spheroid movement was quantified as centroid displacement per dome. Centroid displacement was visualised as spatial heatmaps preserving the physical layout of the array.

### Spheroid Growth Analysis

During the long-term culture of A549 spheroids (up to 35 days), medium change was performed every 2 days, and large images of spheroid array were acquired on Day 0, 2, 10, 20 and 35. After segmentation with SAMJ, spheroid growth was quantified by tracking ROIs within individual dome over time. The primary growth metric presented was total ROI area detected per dome. Roundness of the largest ROI within each dome was used as the representative shape metric.

### Three-dimensional nuclei segmentation

Nuclear instances were segmented in 3D with the StarDist-based DeepStar3D model trained on synthetic volumes^12^, which requires no manual annotation. Stacks were resampled to the training voxel size (0.8 × 0.8 × 1 µm^3^) and processed in [512 × 512] tiles, using probability and non-maximum-suppression thresholds of [0.5] and [0.2].

### Image analysis and quantification

3D reconstruction and image representation were generated using Imaris 10.2 (Bitplane/Oxford Instruments, Zürich, Switzerland) and Fiji (ImageJ). Graphs were generated using Prism 11 (GraphPad Software, San Diego, CA, USA).

## Notes

### Competing Interest Statement

The authors have declared no competing interest.

### Summary of Updates

Author list and acknowledgements section updated.

